# Feed-forward and Feedback Control in Astrocytes for Ca^2+^-based Molecular Communications Nanonetworks

**DOI:** 10.1101/177154

**Authors:** Michael Taynnan Barros, Subhrakanti Dey

## Abstract

Synapses plasticity depends on the gliotransmitters’ concentration in the synaptic channel. And, an abnormal concentration of gliotransmitters is linked to neurodegenerative diseases, including Alzheimer’s, Parkinson’s, and Epilepsy. In this paper, a theoretical investigation of the cause of the abnormal concentration of gliotransmitters and how to achieve its control are presented through a Ca^2+^-signalling-based molecular communications framework. A feed-forward and feedback control technique is used to manipulate IP_3_ values to stabilise the concentration of Ca^2+^ inside the astrocytes. The theoretical analysis of the given model aims i) to stabilize the Ca^2+^ concentration around a particular desired level in order to prevent abnormal gliotransmitters’ concentration (extremely high or low concentration can result in neurodegeneration), ii) to improve the molecular communication performance that utilises Ca^2+^ signalling, and maintain gliotransmitters’ regulation remotely. It shows that the refractory periods from Ca^2+^ can be maintained to lower the noise propagation resulting in smaller time-slots for bit transmission, which can also improve the delay and gain performances. The proposed approach can potentially lead to novel nanomedicine solutions for the treatment of neurodegenerative diseases, where a combination of nanotechnology and gene therapy approaches can be used to elicit the regulated Ca^2+^ signalling in astrocytes, ultimately improving neuronal activity.

## I. Introduction

The field of biological and medical science has witnessed, in recent years, the impact from multidisciplinary research efforts that utilise engineering theories, materials and technologies. Examples of this impact include new approaches for nanomedicine [1], [2], smart drug delivery systems [3] and optogenetics [4], which has seen the fields of nanobiotechnology and information technology being brought together. Telecommunication engineers now are investigating biological communication processes that can either be used to understand the biological signalling processes or develop artificial communication systems at the nanoscale. The latter research topic is known as *Molecular Communication* [5], and its potential applications include sensor and actuator nanonetworks for the human body, as well as new forms of environmental monitoring for smart cities [6].

Inside the human brain, there is a plurality of communication systems with an interplay of different domains such as the electro-chemical synapses, involving multi-scalability in time and space. This phenomenon has attracted massive attention from the recently-formed molecular communication community due to the many communication issues present within neuronal signalling, neuronal networks and cortical circuits. On the other hand, the historically studied bipartite synapse (neuron-to-neuron) is now accepted as the *tripartite synapses*, which are formed by a three-way communication of a pre-synaptic neuron, an astrocyte and a post-synaptic neuron [7] [8]. The internal Ca^2+^ concentration can be influenced by post-synaptic voltage and the communication between astrocytes inside a network of cells and affect the gliotransmitter concentration in the synaptic channel [9]. An example of one gliotransmitter that is triggered by increasing the Ca^2+^ signals in astrocytes [10] is *glutamate*. Through the *Glutamate Dependent NMDA Receptors* (*GNMDAR*), the astrocytes play a significant role in numerous brain processes such as plasticity, learning and memory processes [11]. Therefore stable regulation of Ca^2+^ signalling in astrocytes and their communication is critical. **Communication problems in the tripartite synapses** can lead to serious diseases, including Alzheimer’s, epilepsy, schizophrenia, Parkinson’s and depression [12]. The modelling of these communication processes and the application of control theoretic methods for Ca^2+^ signalling have been previously suggested as a prospect for prevention of neurodegenerative diseases [13].

To this end, a control theoretic model to achieve stable levels of intracellular Ca^2+^ signalling is proposed for deliberately influencing the gliotransmitter concentration indirectly and remotely inside astrocyte communications nanonetworks. A theoretical framework is developed based on a **feed-forward and feedback** control methodology, to maintain stable astrocytes’ cytosolic Ca^2+^ concentration. Since proteins more easily stimulated, IP_3_ is used as a regulation point where its control leads to accurate stimulation of Ca^2+^ ions [14]. This framework opened the door for further contributions including:

- *Regulation of Ca*^2+^ *Signalling in Astrocytes* - A mathematical framework is developed to calculate the desired Ca^2+^ levels based on a given desired IP_3_ value and also the stability of the system (the system in this context is the point of communication between the astrocyte and neuron inter-cellular signalling). Finally, a disturbance analysis is used to investigate the benefits of having a feed-forward component in the control design. <u>The proposed technique achieves the control of Ca^2+^ ion levels in the cytosol of astrocytes with proven stability</u>.
- *Molecular Communication System Design based on Synthetic Biology* - Both transmitter and receiver nanomachines are also designed to incorporate the envisioned synthetic biology circuitry that mimics the control theory model and produces the same output. <u>A toggle switch gene transcription is used to activate control of Ca^2+^ serving as a signal equaliser</u>.
- *Improved Performance of Molecular Communication System* - The use of Ca^2+^ signalling for molecular communication can result in a high quantity of noise propagating through the tissue, resulting in low data rates. However, the control theoretic method proposed in this paper regulates the Ca^2+^ levels, resulting in a minimum amount of noise, which in turn <u>achieves superior communication performance with increased the data rate, increased gain and decreased delay</u>.

The paper is organized as follows. §II introduces the tripartite synapses and the intracellular Ca^2+^ signalling model for astrocytes. §III presents the oscillation behaviour of regular Ca^2+^ signalling process and the problem statement. §IV presents the feed-forward and feedback control technique for astrocytes’ cytosolic Ca^2+^ concentration regulation followed by a stability analysis. §V presents the designed toggle switch biological synthetic circuit. §VI presents the Ca^2+^signallingbased molecular communication system that uses the synthetic circuit to implement the control function. §VII presents the numerical results and analysis of the application of the control technique for elimination of Ca^2+^ signalling oscillations, disturbance, data rate, molecular gain and molecular delay improvements. §VIII presents a discussion about the future envisioned applications. Finally, §IX concludes the paper.

## II. Tripartite Synapses

Fig 1 shows an accurate illustration of the tripartite synapses. The concentration of *gliotransmitters* in the region connecting both the neurons and the astrocyte is crucial for synapses’ plasticity. Researchers have been able to identify the importance of the astrocytes in the tripartite synapses [8], and the communication process is illustrated in Fig 2. In the tripartite synapses, the stimulation of the IP_3_ production in the astrocytes will initiate the Ca^2+^ signalling process. This stimulus starts either from the post-synaptic, pre-synaptic and other astrocytes in the form of gliotransmitters or adenosine triphosphate (ATP). The increased IP_3_ values triggers the release of Ca^2+^ ions into the cytosol from the endoplasmic reticulum. High quantity of Ca^2+^ ions then provoke the release of *glutamate*^1^ into the synaptic channel. Glutamate is also released from the pre-synaptic neuron invoking an increase of Ca^2+^ concentration in the astrocytes. These glutamate molecules go back to the pre-synaptic terminal either to inhibit or assist further glutamate release. Therefore, the intracellular Ca^2+^ signalling in astrocytes of the tripartite synapses dynamically regulates synaptic transmission. It is still debatable also the influence of different types of neurons on the signalling in astrocytes, and the absence of the modulatory activity from pre-synaptic neurons and interneurons [15]. Therefore, we initially neglect these effects for the sake of defining an initial scenario whereby control can be achieved. Based on this, the focus is on the core of this sequential communication process by concentrating on the intracellular Ca^2+^ signalling process in astrocytes.

**Fig. 1:**
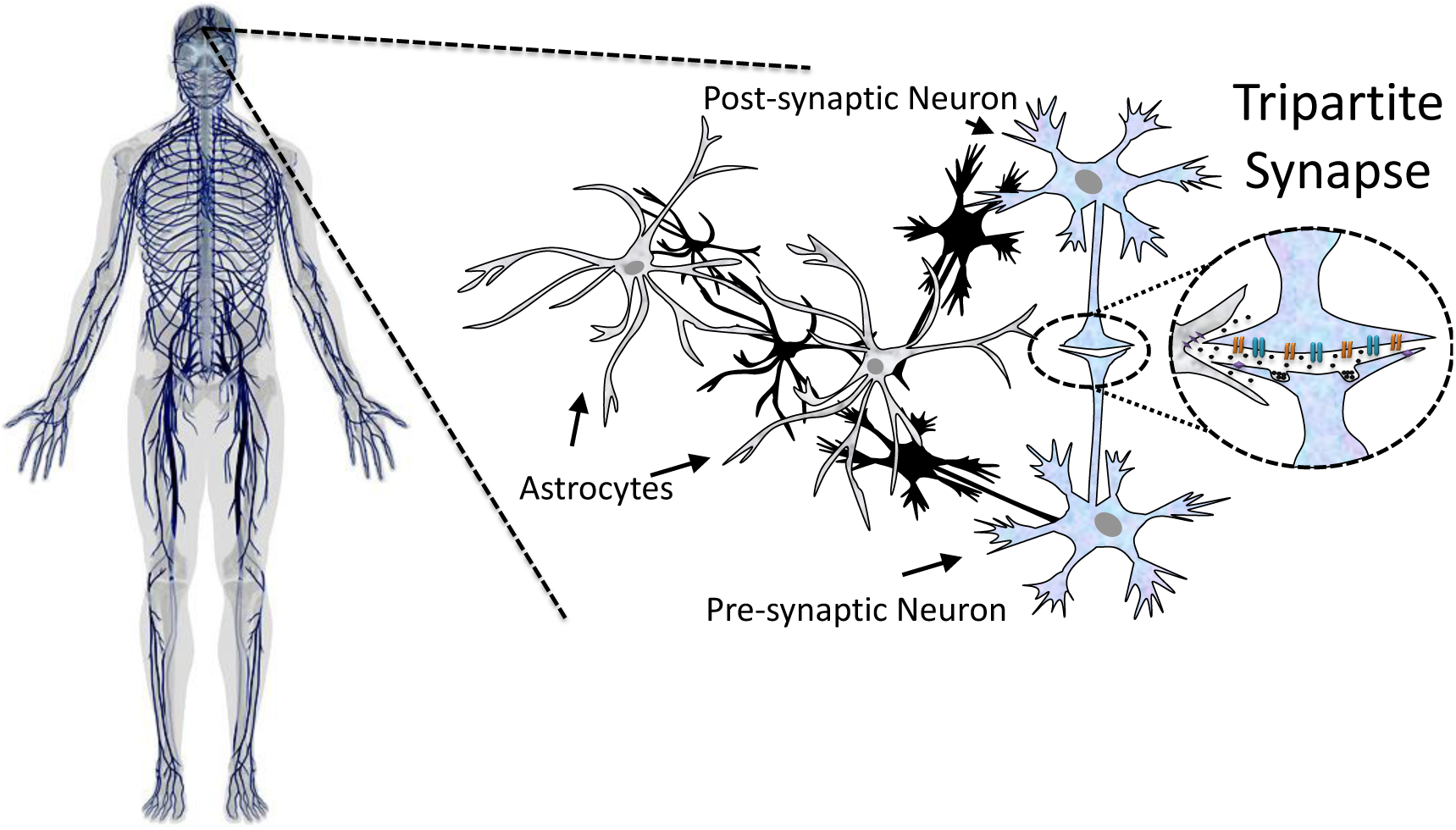
Tripartite synapses overview, showing the communication process between the astrocyte cell and the pre-synaptic neuron, as well as the post-synaptic neuron. The three-way communication process emits *gliotranmitters* which are molecules crucial for maintaining synaptic transmission quality.

**Fig. 2:**
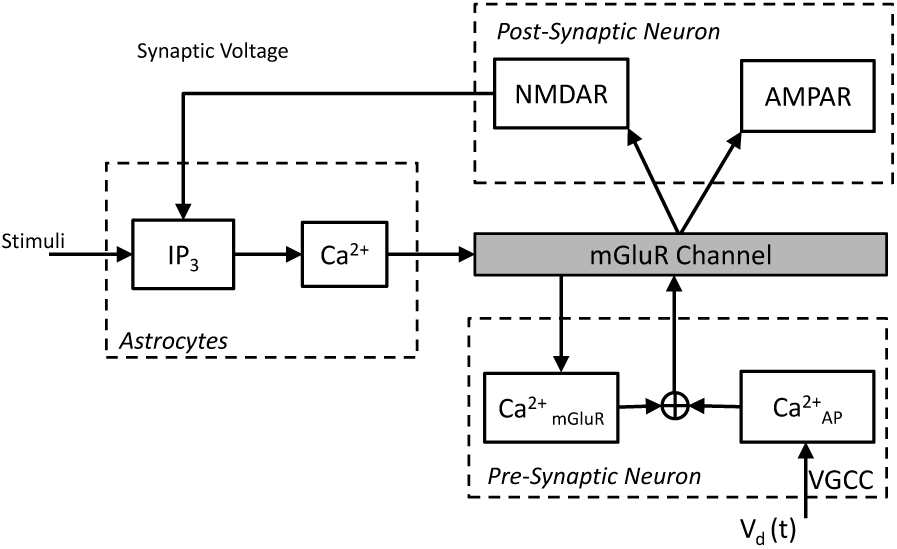
Model for the Tripartite Synapses. Pre-synaptic neuron, postsynaptic neuron and the astrocyte communicates through a gliotransmitter channel that is invoked from Ca^2+^ signalling.

The intracellular Ca^2+^ signalling model in astrocytes consists of state equations for the Ca^2+^ concentration in the cytosol (*C*) (Eq. 1), kinetics of IP_3_ receptors (*h*) (Eq. 2) as well as the IP_3_ concentration (*I*) (Eq. 3). This model is proposed in [16], and a visual illustration of the model is presented in Fig. 3. The choice of this model is twofold i) needs to be simple and accurate enough to create a low complexity control model ii) it is validated with experiments of astrocytes. The main state equations are defined as follows:

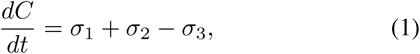

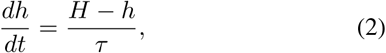

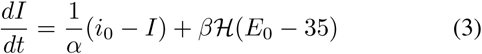

where *α* is the constant degradation time of IP_3_ concentration, *i*_0_ is the IP_3_ concentration in equilibrium, *β* is the production rate of IP_3_ ions, *E*_0_ is the pre-synaptic potential and ℋ (.) is the Heaviside function. *Ca*^2+^*-induced Ca*^2+^ *release* (*CICR*) is the trigger process of Ca^2+^ ions from the *sarco(endo)plasmic reticulum* by existing Ca^2+^ ions within the cytosol. The quantities *H,τ* are defined in (8), (9, (10). The function *σ*_1_ models the *CICR* and is defined as:

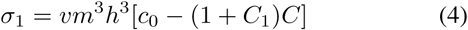

where *v* is the maximal *CICR* rate, *c*_0_ is the total cell free Ca^2+^ concentration depending on the cytosol volume, and *C*_1_ is the ratio between the cytosol and endoplasmic reticulum volume.

**Fig. 3:**
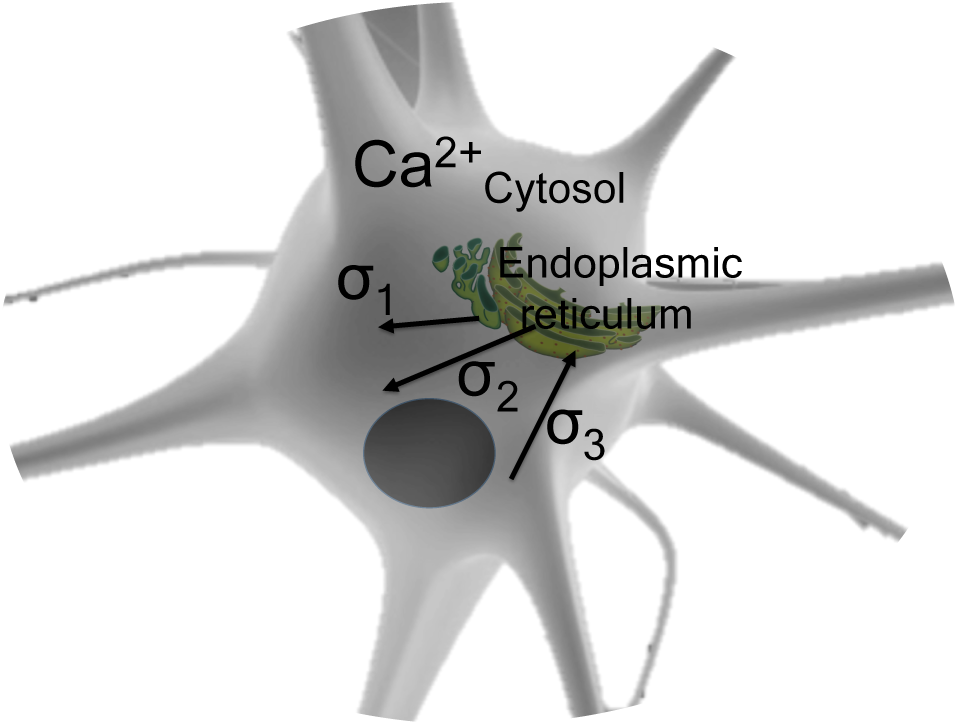
Intracellular Ca^2+^ signalling model for astrocytes. The flux/efflux rates control the concentration of Ca^2+^ signalling. The function *σ*_1_ models the Ca^2+^-induced Ca^2+^ release (*CICR*). The function *σ*_2_ denotes the leakage of Ca^2+^ ions from the *endoplasmic reticum* to the cytosol, and the function *σ*_3_ denotes denotes the SERCA-driven Ca^2+^ uptake from the cytosol into the *endoplasmic reticulum*.

The IP_3_ and Ca^2+^ ion binding process that is responsible for providing stable IP_3_ kinetics is represented as:

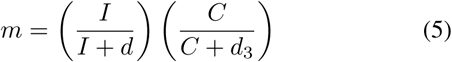

where *d* is the IP_3_ dissociation constant and *d*_3_ is the Ca^2+^ activation-dissociation constant.

The function *σ*_2_ denotes the leakage of Ca^2+^ ions to the cytosol from the *sarco(endo)plasmic reticum* and is represented as:

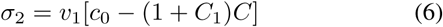

where *v*_1_ is the maximal rate of Ca^2+^ ions leakage from the endoplasmic reticulum.

The efflux of Ca^2+^ from the *sarco(endo)plasmic reticulum* to the *endoplasmic reticulum* (*SERCA*) is represented as:

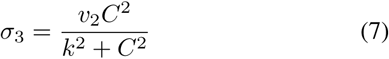

where *v*_2_ is the maximal rate of SERCA uptake, and *k* is Ca^2+^ binding affinity.

The following equations are important for modelling *h*:

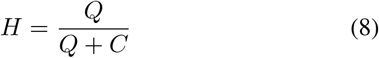

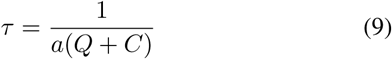

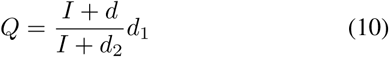

where *d*_1_ is the Ca^2+^ inactivation dissociation constant, *d*_2_ is the IP_3_ dissociation constant and *a* is the IP_3_ receptors binding rate for Ca^2+^ inhibition.

## III. Problem Statement

Neurodegenerative diseases are related to the quality of the synapses (plasticity) in neuronal communication. The poor concentration of glutamate inside the synaptic channel will lead to poor propagation in the synapses, causing lack of memory, insomnia, depression which are symptoms of most neurodegenerative diseases. Current treatment techniques of neurodegenerative diseases are based on drugs that are not efficient and only helps to eliminate symptoms and not treat them. Based on what has been already presented, achieving Ca^2+^ control in astrocytes can indirectly control the glutamate release and potentially improve the synaptic plasticity. Control of Ca^2+^ has been theoretically achieved in our previous work [17]. The primary challenge now is to provide an analysis on the astrocytes Ca^2+^ concentration of the tripartite synapses and, therefore, create a theoretical framework that supports the indirect control of glutamate.

For this, the problem of controlling levels of Ca^2+^ in the cytosol is investigated in this paper and its influence on an astrocyte network communication. More specifically, internal Ca^2+^ signalling is characterised by oscillations invoked by a particular range of IP_3_. Fig 4 shows the Ca^2+^ oscillation at IP_3_ = 0.5 *μ*M and when production rate of IP_3_ ions (*β* - Eqn. 3) varies from 0.1-1.5 *μ*M/s and modulates both the amplitude and frequency of the signal.

**Fig. 4:**
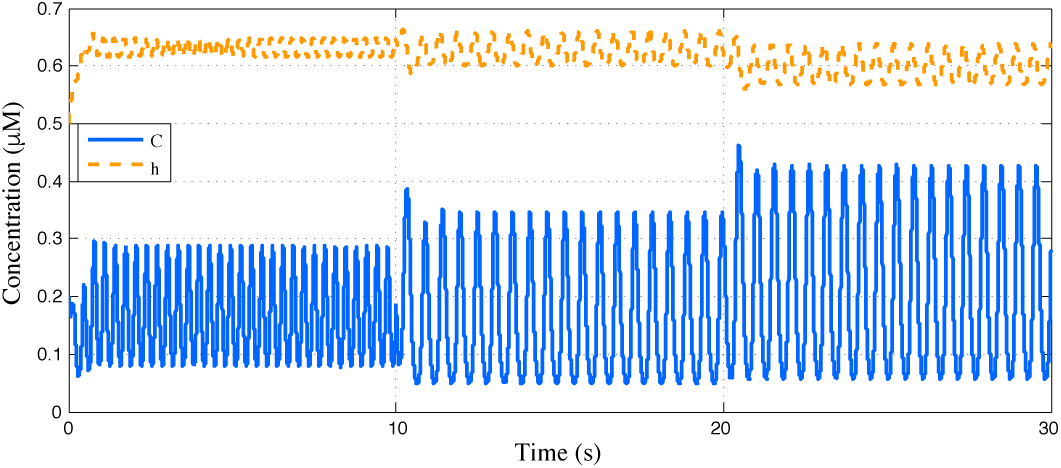
Ca^2+^ oscillation with respect to time. In this illustration the IP_3_ = 0.5 *μ*M. The Ca^2+^ concentration (*C* - blue line) oscillates alongside with the kinetics of IP_3_ receptors (*h* - dashed yellow line).

Fig 5 shows how the IP_3_ can affect the intracellular Ca^2+^ signalling based on the postsynaptic voltage influence. An increase of IP_3_ in the system is desired for regular Ca^2+^ concentration levels. Since *β* is responsible for the IP_3_ increase, it is highly critical for regulation of Ca^2+^ concentration levels. As soon as IP_3_ achieves a constant level, the Ca^2+^ concentration will drop. The post-synaptic voltage will also be influenced, as seen in Fig. 5 (*E*_0_ = 50*V* in the left, *E*_0_ = 35*V* in the center and *E*_0_ = 0*V* in the right). However, since the synapses happen periodically, we assume only activated neurons and also no inter-synaptic interference in the synaptic channel. The IP_3_ is, in conclusion, a decisive factor for Ca^2+^ regulation, in which its increase is controlled by *β*. We further explore it with a mathematical model that enables the intracellular Ca^2+^ signalling control by regulating IP_3_ levels.

**Fig. 5:**
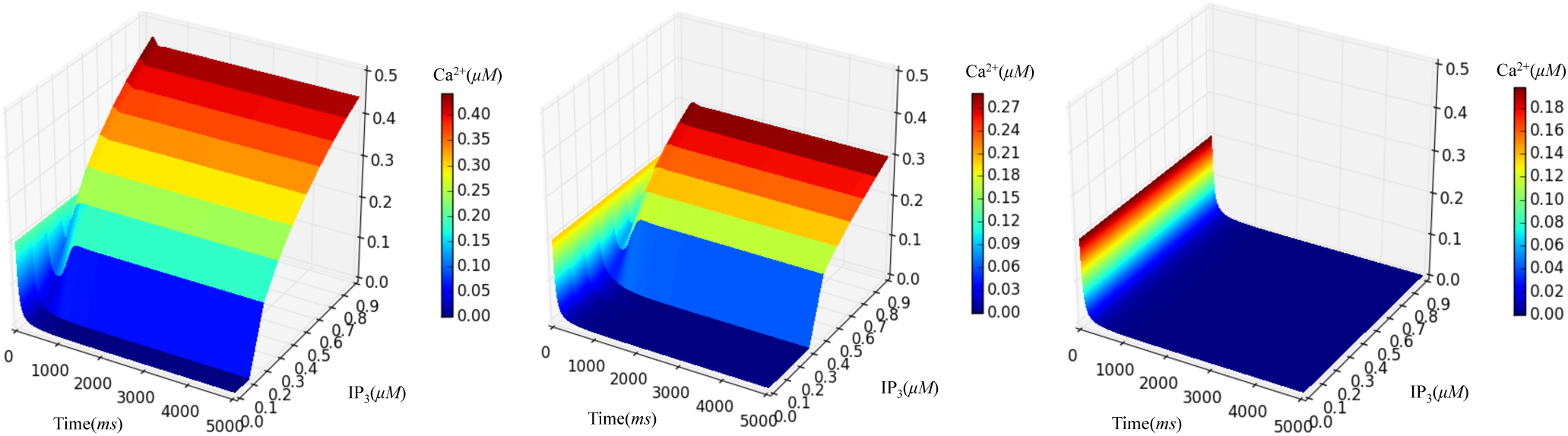
Effect of both IP_3_ and the *E*_0_ on the Ca^2+^ concentration, *E*_0_ = 50*V* in the left, *E*_0_ = 35*V* in the center and *E*_0_ = 0*V* in the right. The Ca^2+^ concentration is highly dependent on the increasing factor of IP_3_ and, therefore, so is the stability.

## IV. Feed-forward and Feedback Control of Intracellular Ca^2+^ Signalling in Astrocytes

Since IP_3_ activation can only be performed in highly controlled settings [17], its regulation of Ca^2+^ concentration is more suitable for *in-vitro* scenarios at the moment. The proposed approach can be potentially realised with the help of nanoparticles or silicon devices inside a cell to inhibit or activate Ca^2+^ as part of the control model.

Fig. 6 shows a functional block diagram of the desired Ca^2+^ signalling set point regulation. Before proceeding further, assume for the rest of the paper that ℋ(*E*_0_ − 35) = 1. Note that for other values of the Heaviside function (such as 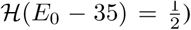, a similar analysis can be applied. By denoting the column vector state **x** = [*C h I*]’, rewrite (1), (2), (3) as the following nonlienar controlled state space system

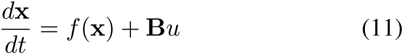

where *f*(x) = [*f*_1_(x) *f*_2_(x) *f*_3_(x)]′ and *f*_1_(.) = σ_1_+σ_2_+σ_3_,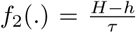, and 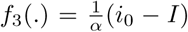 and **B** = [0 0 1]′. The control variable *u* represents the state feedback and feedforward based IP_3_ regulation parameter given by (see also [14]):

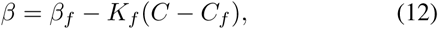

where *β*_*f*_ is the feed-forward control representing the desired IP_3_ level *I*_*f*_, *C*_*f*_ is the associated desired Ca^2+^ concentration level and *K*_*f*_ is the linear feedback gain. Note that although not visible in the above equation, there is an associated value of *h* as well, which is denoted by *h*_*f*_. Denote the entire 0 associated state vector as **x**_**f**_ = [*C*_*f*_ *h*_*f*_ *I*_*f*_]′. Then it follows that

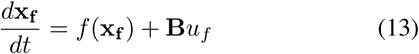

where of course, *u*_*f*_ = *β*_*f*_. Here, the measured output is the Ca^2+^ concentration level, so that at the desired level, the output is given by 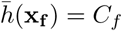.

*Remark 1:* Note that *without loss of generality*, one can assume that in the uncontrolled case, i.e., when *β* = 0, there is an equilibrium point at the origin for the nonlinear dynamical system 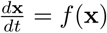 [14], implying *f*(**0**) = 0. This fact will be used in the stability analysis of the system when the control law (12) is applied in a subsequent subsection.

**Fig. 6:**
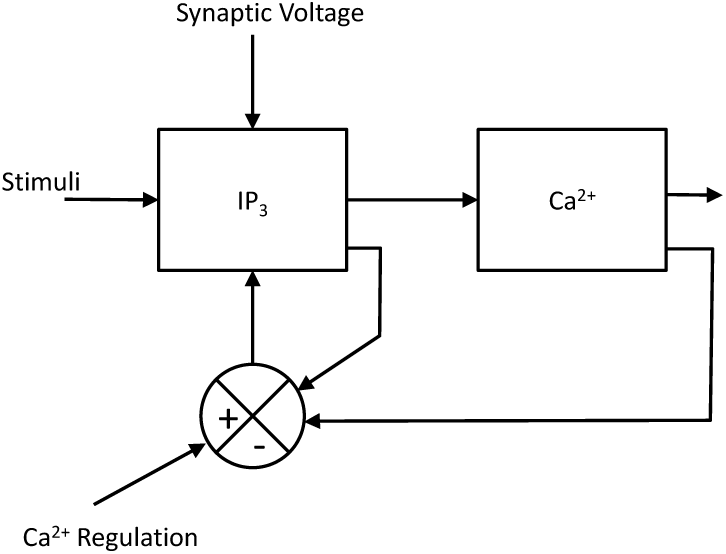
Ca^2+^ regulation flowchart based on the regulation of cytosolic IP_3_. The input rate of IP_3_ is a function of the current cytosolic Ca^2+^ and the current IP_3_ levels, which makes it both a state feedback and feed-forward control.

A similar control law to (12) *including the feed-forward term* was proposed for disturbance rejection in roll-to-roll manufacturing system [14]. In this model, no specific disturbance variables are implemented, which may be present in cases of diseases that may affect the normal regulated Ca^2+^ concentration levels. However, since the nature of the problem investigated in this paper is elimination of Ca^2+^ oscillations, it is clear that not only such a control function perfectly fits into our modelling of the nonlinear controlled Ca^2+^ regulation system, but also allows us the flexibility of including disturbance factors in future modelling extensions.

### A. β_f_ and C_f_ Relationship

To obtain the required regulation factor *β*_*f*_ for a desired *C*_*f*_, a mathematical relationship between *β*_*f*_ and *C*_*f*_ is obtained as follows. Suppose there is an equilibrium point (in the controlled case when *β >* 0) at (*h*º *c*º *I*_0_), then 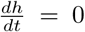 at *h* = *h*º and 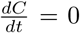 at *C* = *c*º, and 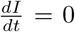 for *I* = *I*_0_. Rewriting Eq. 1 and Eq. 2, obtaining:

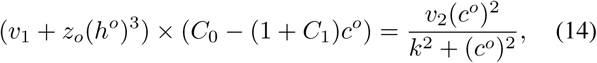

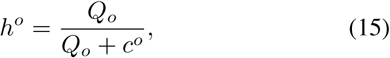

where

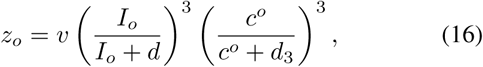

and

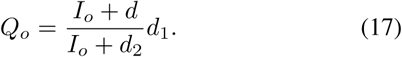

Here *I*_*o*_ is obtained from for Eq. 3, (since ℋ(*E*_0_ − 35) = 1 as previously assumed), by solving:

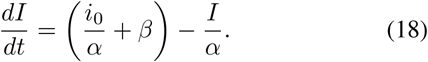

which yields

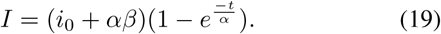

As *t* → ∞, *I* becomes a constant *I*_*o*_, which is represented as:

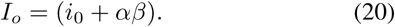

Substituting the above value of *I*_*o*_ in the equations for *Q*_*o*_*,h*º and *z*_*o*_, one can solve the nonlinear equation (14) to obtain *c*º = *C*_*f*_ for a given *I*_*o*_ = *β*_*f*_.

### B. System Stability

Since anomalies in the dynamics in Ca^2+^ signalling can lead to diseases and also tissue death, a stabilising control model for such a system is critically required. In the following, a stability analysis is presented that ensures that how the feed-forward and feedback based regulation of the IP_3_ level can maintain the Ca^2+^ level very close to its desired value. This requires a linearization of the nonlinear dynamics around the equilibrium using Taylor’s series expansion, assuming one initializes the system very near the origin. Recall from Remark 1 that is assumed without loss of generality that there is an equilibrium at the origin for the uncontrolled model.

Note that by carrying out a Talyor’s expansion of (11) around the origin, and ignoring the higher order nonlinear terms (since the expansion is obtained very close to the origin) one obtains

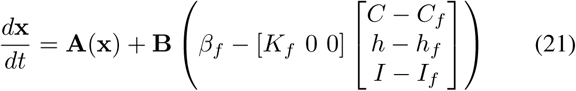

where **Ax** is the linear part of *f*(x). Note that **A** is given by

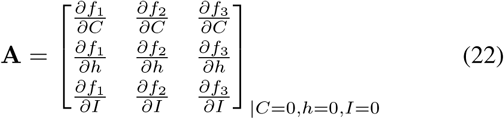

It follows easily that

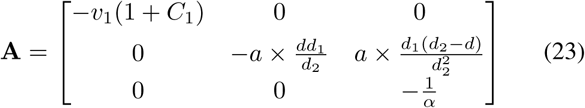

One can also show from linearization of (13) around the origin that the following is true:

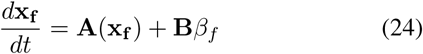

Defining *ζ* = **x** – **x**_**f**_, having (by subtracting (24) from (24))

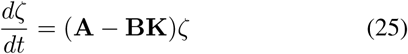

where **K** = [*K*_*f*_ 0 0]. It follows then that the matrix **A**–**BK** is given by

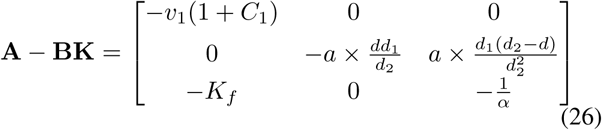

Finally, the eigenvalues of **A**–**BK** can be easily shown to be 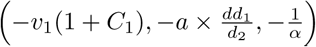. Since all of these eigenvalues are negative, the matrix **A** – **BK** is Hurwitz, which implies that **x**(*t*)-**x**_**f**_(*t*) → 0 as *t* → ∞, or *C* → *C*_*f*_ as *t* → ∞. This proves that under the feed-forward and feedback control law proposed in this work, the Ca^2+^ level can be maintained at the desired level after a sufficiently large time interval. Simulation studies will show that the convergence of *C* to the desired value *C*_*f*_ is considerably fast.

## V. Synthetic Circuit for Control Implementation

The implementation of the control function (12) in astrocytes requires a high-level synthetic circuit design. Synthetic biology has been extensively studied in mammalian cells [18], [19], [20]. However, since the objective of the remainder of the paper is for the validation of the proposed technique, in the following one must find an abstraction of the control function regarding the synthetic circuit. Therefore the implementation of the control function can be expected to be partially achieved in astrocytes. The technique consists of an envisioned programmable genetic circuit that triggers the production of IP_3_, and therefore, controls *Ca*^2+^ through a toggle switch. The advantage of this approach is that there is no need for rewiring *Ca*^2+^ signalling pathways. Previously, RNAi has also been used to screen the function of different players in the *Ca*^2+^ signalling pathway [21], and the *Ca*^2+^ pathway rewiring has been targeted by synthetic biologists [22].

The theoretical two-gene bistable toggle switch has been proposed by Gardner et al [23]. The switch consists of two constitutive promoters (coloured black and grey) and two repressor genes (also coloured black and grey). The black repressor protein silences the grey promoter, which drives production of the grey repressor protein. Conversely, the grey repressor protein silences the black promoter, which drives production of the black repressor protein. Thus, if the black repressor protein were produced, the grey repressor protein could not be produced, and vice versa. This is an example of a synthetic bistable toggle switch similar to the theoretical one in Figure 7. The presented approach has the goal to only theoretically simulate an dependent stimuli of IP_3_ production based on current Ca^2+^ levels.

**Fig. 7:**
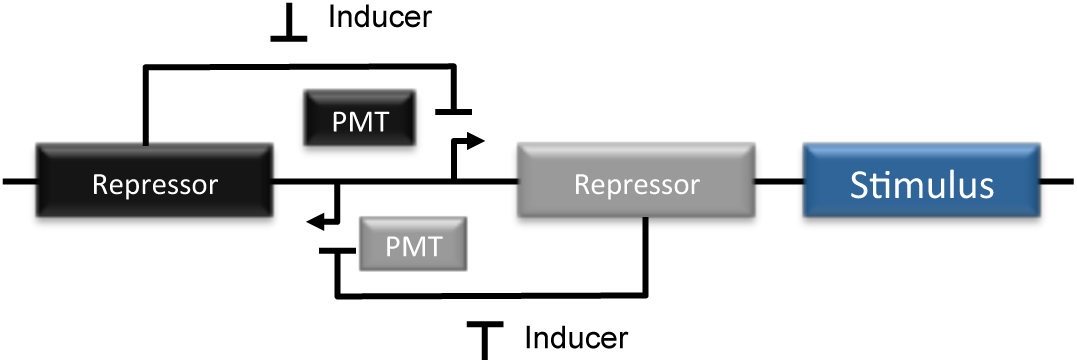
The black gene “repressor” is transcribed from its black promoter (PMT). The black repressor protein binds to the grey promoter to block the production of the grey repressor protein. The grey repressor protein blocks the production of the black repressor protein when the grey repressor protein binds to the black promoter. To detect which state the toggle switch is in, Stimulus was placed downstream of the grey promoter so that the grey repressor and stimulus are produced simultaneously. Once stimulus is produced, the IP_3_ production starts as well.

The control function (12) requires monitoring of both *Ca*^2+^ and IP_3_ states. This can be achieved by individual cells simultaneously transfected with recombinant, *Ca*^2+^-sensitive and IP_3_-sensitive fluorescent reporters [24], [25]. This is abstracted here with the assumption of ideal monitoring of both *Ca*^2+^ and IP_3_ levels since the design of the synthetic circuitry is not the focus of the paper.

### A. Stochastic Solver

## VI. CA^2+^-signalling-based Molecular Communication System Model

In this section, a single mathematical framework merges the *Ca*^2+^-signalling, control and synthetic biology models alongside diffusion and 3D cellular tissue modelling. Barros et. al. has extensively studied this model [26], [27], [28].

Spatio-temporal dynamics is captured by a 3D cellular tissue model, correlating phenomena and variables at different scales and analysing how the control technique will perform. The description is composed of two parts: the 3D modelling of the cellular tissues, and the a stochastic model for the scheduling of reactions within individual cells.

### A. 3D Modelling of Cellular Tissue Structure

Consider a cellular tissue space (*S*) composed of *I* × *J* × *K* cells (*c*), where *c*_*i,j,k*_ (*i* = 1*…I*; *j* = 1*,…J* and *k* = 1*,…K*) denotes an arbitrary cell in the tissue. The cells are connected with a maximum of six neighbouring cells. The topological organisation of the astrocytes is considered following regular specifications in [29]. A Stochastic Solver computes the values of each pool over time, selecting and executing scheduled reactions. The pool is negatively or positively affected by a constant *α* when a particular reaction is run. More details on the astrocytes’ pools and reactions can be found in Section II.

Modelling diffusion in a cellular tissue area captures the temporal-spatial dynamics of intercellular Ca^2+^ signaling. The model considers Ca^2+^ concentration difference for temporal-spatial characteristic as follows:

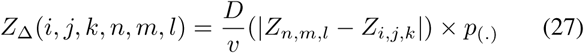

where *n* ∈ (*i*−1*,i*+1), *m* ∈ (*j*−1*,j*+1), *l* ∈ (*k*−1*,k*+1), *D* is the diffusion coefficient, *v* is the volume of the cell, and *Z*_Δ_ is the difference in Ca^2+^ concentration between the cells. *p*_(.)_ is the probability of the gap junction opening. More details of this model can be found in [27].

At each time step, the Gillespie algorithm [30] (a Stochastic Solver) schedules a particular astrocyte ∈ *S*, its reaction (*R*) and the time duration of this particular reaction (*t*), therefore determining the quantity of each pool over time. Each reaction is allocated a reaction constant (*a*_*r*_). Considering that *α*_0_ is the summation of all *a*_*r*_ in *R*, the next reaction chosen *r*_*u*_ will be:

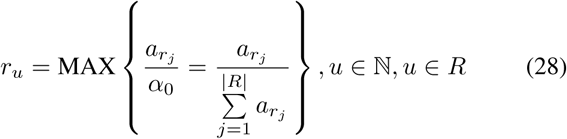

which follows the *roulette wheel selection* process, by choosing the events based on their probability values. However, *u* must satisfy the following restriction:

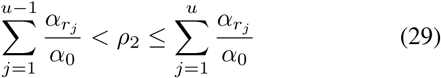

in which *ρ*_2_ is a uniform random variable with values in the range (0,1).

At each time step (*t*), a time lapse (*τ*_*t*_) is derived based on *α*_0_, and is represented as:

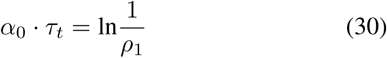

in which *ρ*_1_ is a uniform random variable with values in the range (0,1). This process ends when 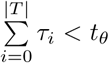, where *T* is the set of *t* and *t*_*θ*_ is the maximum simulation time.

### B. System Design

Both transmitter and receiver design are based on [26], [27], [31] plus the following considerations. Both transmitter and receiver have been engineered with the proposed control function and its toggle switch activation in the form of an equaliser. In this way, both devices are capable of ideally responding to a desired level of IP_3_ and, therefore, maintain stable levels of Ca^2+^ concentration during the transmission period *T*_*b*_. The toggle switch is activated with the two following processes: 1) the external cellular stimuli in the transmitter and 2) the upcoming Ca^2+^ signals in the receiver. This process is depicted in Fig. 8. The system design also has the following assumptions. *Assumption 1:* Synchronization between the transmitter and receiver is considered in ideal settings [26]. *Assumption 2:* The effects of the synthetic circuit on the stimulus of IP_3_ is also considered ideal. The paper only concentrates on the effect of the Ca^2+^ control in the astrocyte communication. *Assumption 3:* The used modulation is OnOff Shifting Keying. We do not extend the analysis for other types of modulation.

**Fig. 8:**
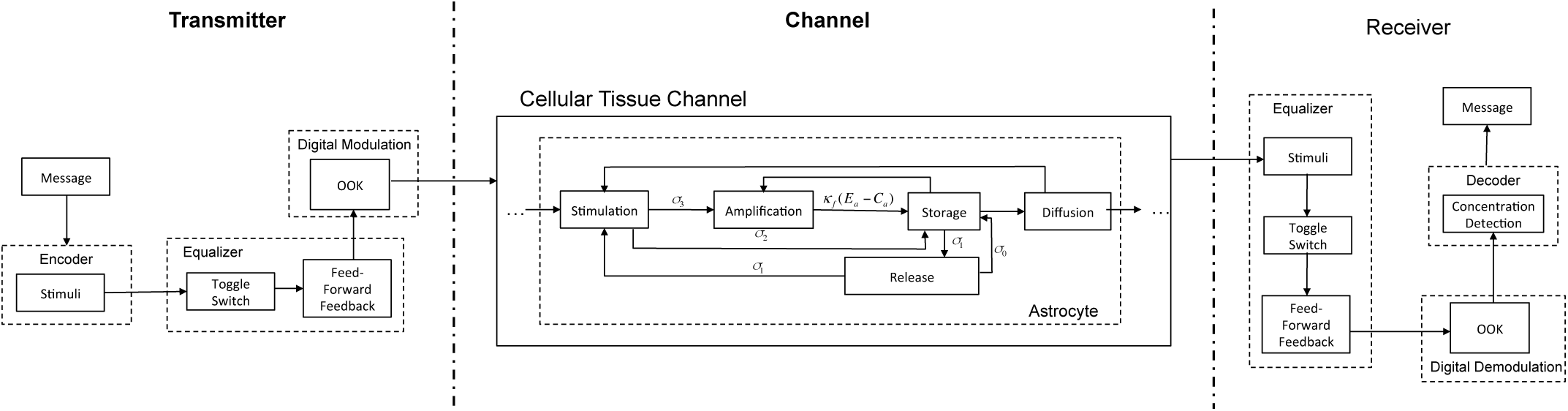
System design. The transmitter has an equalizer to inhibit intracellular noise in the generated cellular signalling. The channel is represented by a cellular tissue. The receiver is also equipped with an equalizer to regenerate the signal affect by noise in different levels.

## VII. Performance Analysis

The performance analysis of the proposed regulation of Ca^2+^ concentration levels for astrocytes is in the following. It is divided into four parts for a proper understanding of the system under controllable conditions. First, how the control system eliminates intracellular Ca^2+^ oscillations in astrocytes is shown by solving the problem defined in Section III. This is followed by the disturbance analysis, where Gaussian noise is applied to the intracellular Ca^2+^ signalling for adding a controlled abnormal behaviour to the system and observing system effectiveness while looking at the feed-forward and feedback techniques separately. Finally, analyses of the performance improvement for the Ca^2+^-signalling-based molecular communication system of astrocyte networks is shown.

*1) Elimination of Intracellular Ca*^2+^ *Oscillations:* As mentioned in Section III, the elimination of the intracellular Ca^2+^ oscillation is a desired outcome of the control process. For this Eqs. 1, 2 and 3 are solved using the parameter values found in Table I. For the control technique, the *β* in Eq. 3 is replaced by Eq. 12, in order to integrate the feed-forward and feedback control element to the system. A value of *C*_*f*_ = 0.32 *μM* is chosen, which is a central value of the system, and leads to the desired value of *β*_*f*_ with an appropriate calibration of the *K*_*f*_.

A total elimination of the Ca^2+^ oscillation is obtained using the proposed mechanism, and this is illustrated in Fig 10. Eq. 12, which represents the state feedback and feed-forward control, can efficiently adjust *β* accordingly and maintain Ca^2+^ concentration levels throughout the period shown across all Ca^2+^ signal modulation variations. This positive result demonstrates the effectiveness and potential of utilising the control technique to stabilise the excessive Ca^2+^ concentration that may either lead to neurodegenerative diseases or artificial molecular communication, which will be further explored in the following subsections. Further results with a different set of scenarios are presented in [17].

*2) Analysis of System Disturbance:* Feed-forward control techniques are used in control theory to stabilise systems under the presence of disturbances. The presented Ca^2+^ signalling model so far does not include any disturbances or noise components. To analyse the benefit of the feedback control technique with and without feed-forward separately, a noise component is integrated and determine the effectiveness of the approach to stabilising the disturbance. To achieve this, a disturbance component was added to Eq. 1, through addition of Gaussian noise [32]. This additional noise affects the Ca^2+^ level in the astrocyte cell’s cytosol.

**Fig. 9:**
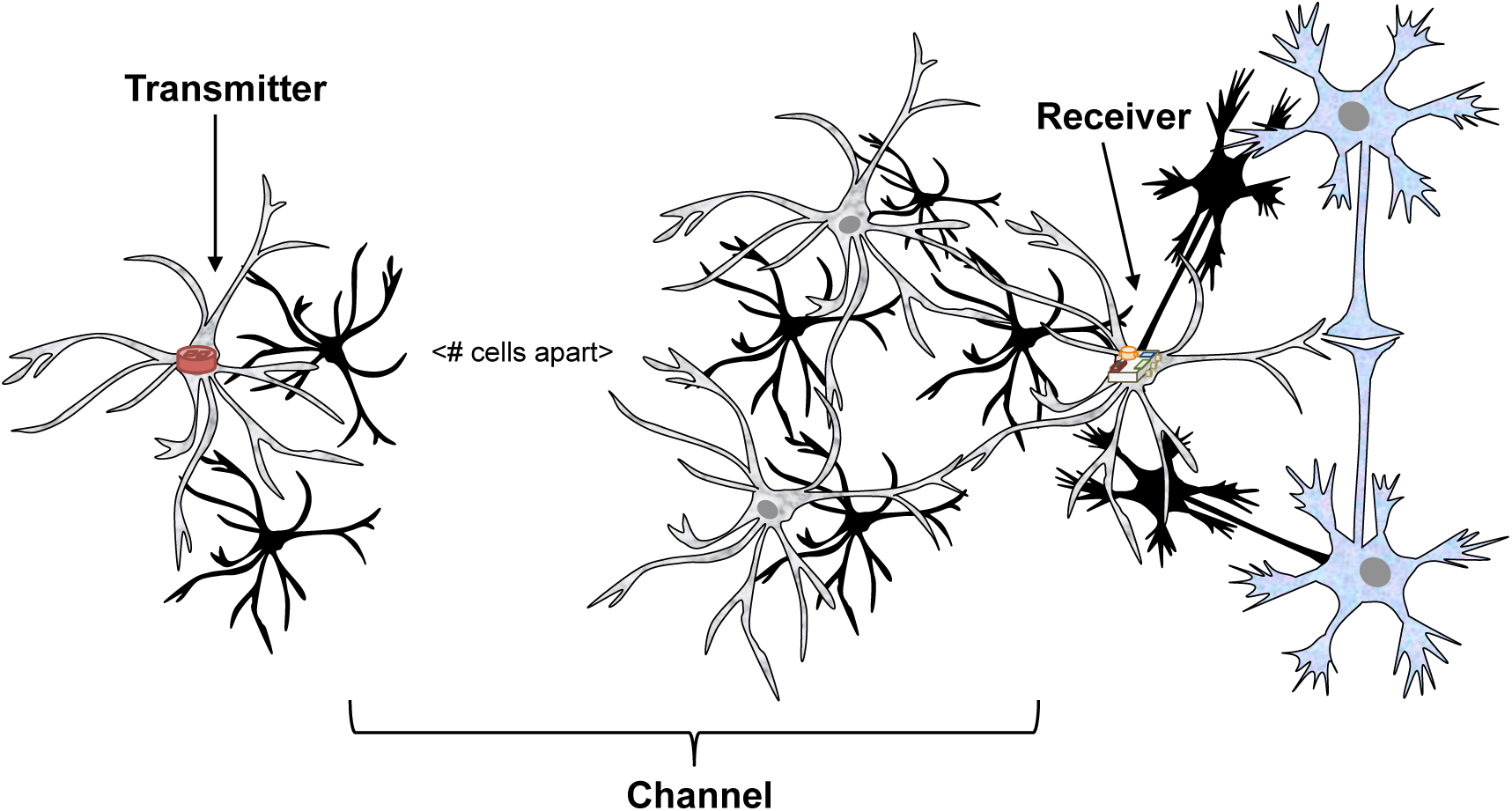
Astrocyte network communication system. The synthetic transmitter and receiver are placed inside a astrocyte cellular tissue. However, the receiver is always in interface with other two neurons forming a tripartite synapses.

**TABLE I.** Simulation parameters for astrocytes.

| Variable | Value |
| --- | --- |
| $v$ | $6s^{-1}$ |
| $v_1$ | $0.11 s^{-1}$ |
| $c_0$ | $2.0\mu M$ |
| $C_1$ | 0.185 |
| $v_2$ | $0.9 Ms^{-1}$ |
| $k$ | $0.1\mu M$ |
| $d$ | $0.13\mu M$ |
| $d_1$ | $1.049 s^{-1}$ |
| $d_2$ | $0.9434\mu M/s$ |
| $d_3$ | $0.08234\mu M$ |
| $a$ | $0.2 s^{-1}$ |
| $\alpha$ | $1/0.00014\mu M$ |
| $i_0$ | $0.160\mu M$ |
| $\beta$ | $0.1-1.5\mu M$ |
| $E_0$ | 35 |

**Fig. 10:**
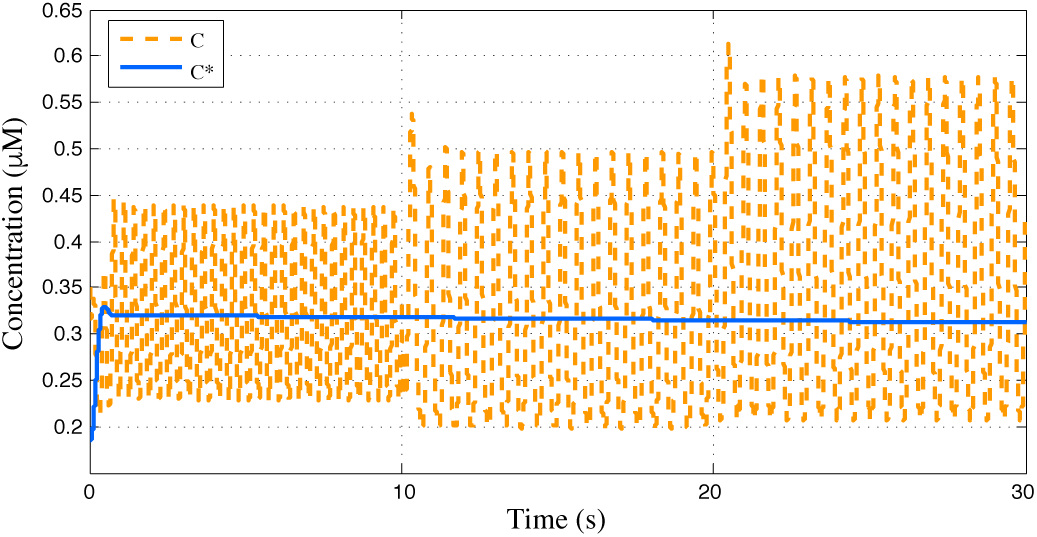
Elimination of Ca^2+^ oscillation using the proposed feedback and feed-forward control technique. Regular Ca^2+^ oscillations *C* (dashed line) is compared to a controlled Ca^2+^ level *C*^*^ (straight line) for IP_3_ = 0.5 (*μM*).

As illustrated in Fig. 11, the feed-forward control can maintain the levels of Ca^2+^ even with the additional Gaussian noise. This is obtained by the relation of the current Ca^2+^ concentration levels with the desired level, and therefore, this stability is achieved through the adjustment of *β*. This demonstrates the excessive control fluctuations in Ca^2+^ concentration levels from additional noise.

**Fig. 11:**
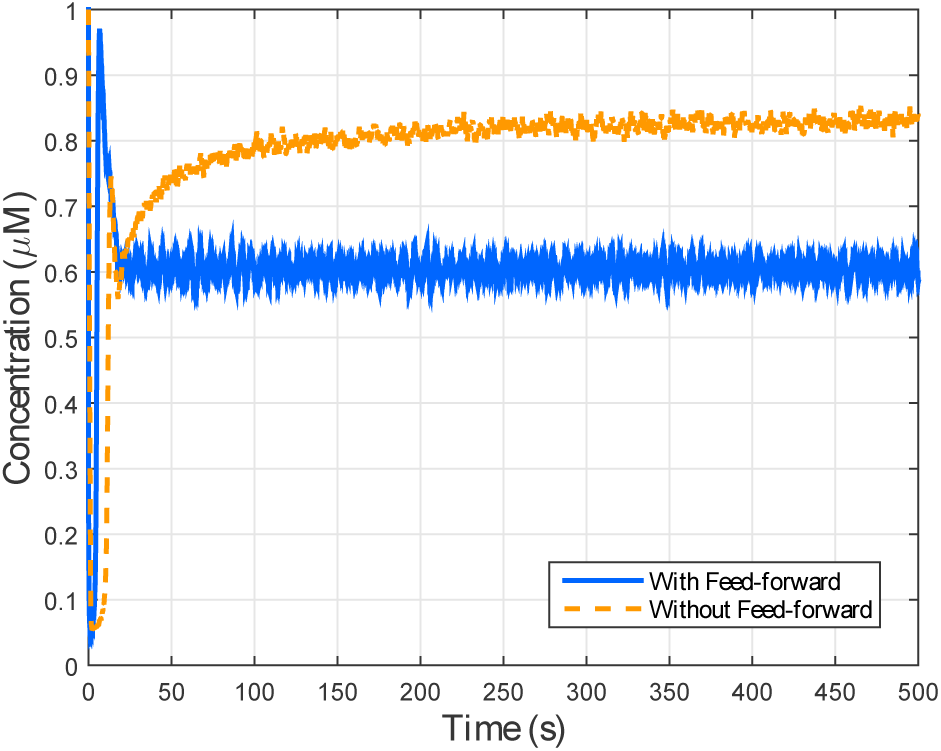
The application of the Feed-forward technique to maintain the desired levels of Ca^2+^ in the astrocytes’ cytosol.

*3) Maintaining Stable Ca*^2+^ *Concentration:* While the previous section presented the case of maintaining stable Ca^2+^ concentration due to excessive noise, in this section, the impact of overall fluctuations in the cytosolic concentration of the cell is discussed. The ability to maintain the stability of the cytosolic Ca^2+^ concentration levels in astrocyte cells can lead to not only a healthy state of the cell but also the synaptic transmission quality in the tripartite synapses. In the event of fluctuations (extreme low or high) in the overall concentration, this can lead to some neurodegenerative diseases. For example, low Ca^2+^ concentration leads to cell death and poor functioning of neurons that cause depression, whereas high Ca^2+^ concentration is linked with one of the causes of Alzheimer’s disease [33], [34], [35]. For evaluation of both extreme high and low concentrations, a simple model is presented that shows the effectiveness of the control technique in maintaining the stable zones of Ca^2+^ concentration levels. We do not claim that this technique can directly be applied to the treatment of neurodegenerative diseases. This method is used only in micro-levels of the astrocyte communication and tripartite synapses. Since neurodegeneration affects higher levels of communication in the brain, there needs to be a extended version of this system that achieves the very challenging distributed astrocyte control.

There is a variation of *β* in Eq. 3 from 0.1 to 0.9 *μM* that is compared with the resulting Ca^2+^ concentration levels with the feed-forward and feedback control technique. In the case of when no control is applied, Ca^2+^ oscillations are expected. Based on the oscillations, the maximum and minimum values are selected of the final concentrations and used them to define three regions: *extreme high region*, *extreme low region*, and *stable region*. The extreme high region consists of any value higher than the maximum Ca^2+^ concentration levels, while the extreme low region contains any value lower than the minimum Ca^2+^ concentration levels. The stable region, which represents the safe level in the Ca^2+^ concentration, is in between the extreme limits.

Regulation of Ca^2+^ concentration can maintain stability in the concentration within the safe region for all IP_3_ values as illustrated in Fig. 12. The *C*_*f*_ from Eq. 12 was selected based on the central value between the maximum and minimum values of *C*. The *β*_*f*_ was computed based on *C*_*f*_, and by adjusting this, the system will change *C* to match with *C*_*f*_. There is a concern of long term usage of the proposed technique in regards to cell function homoeostasis. Even though the Ca^2+^ concentration levels can be maintained for long periods of time (minutes), we do not guarantee that cellular function will not be changed in longer periods. This will require novel protocol solutions that activate and deactivate the control autonomously under certain condition and can be realized through synthetic biology.

**Fig. 12:**
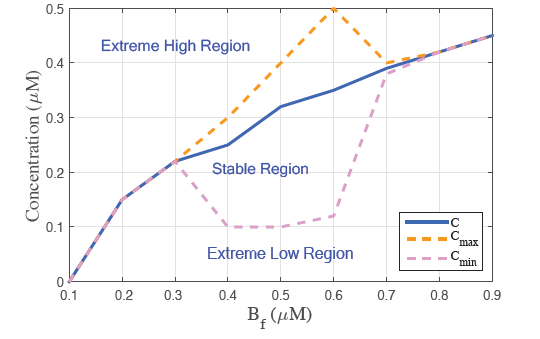
Application of the control model to achieve the desired Ca^2+^ level compared to regions that can result in diseases. *Extreme high region* is any value higher than the maximum Ca^2+^ levels, *Extreme low region* is any value lower than the minimum Ca^2+^ levels, and the *stable region* is the Ca^2+^ levels in between the maximum and minimum values.

*4) Data Rate:* One of the issues regarding the use of Ca^2+^ signalling for molecular communication is the artificial stimulation of the ions for communication purpose, as well as the excessive noise that can result in poor data rate performance. Barros et. al. [26], [27], showed that large *Tb* is desired to achieve reasonable communication capacity but this results in poor data rate for most types of cells that communicate using Ca^2+^ ions. Another reason for the long bit transmission periods *Tb* is due to the refractory time of the Ca^2+^ oscillation. The refractory time is an inherent process found in Ca^2+^ intracellular signalling, where the concentration of the fluctuating ions are required to stabilise before they can be stimulated again, Fig. 13. The objective now is to integrate the proposed feed-forward and feedback control technique towards the refractory time elimination using an equaliser.

**Fig. 13:**
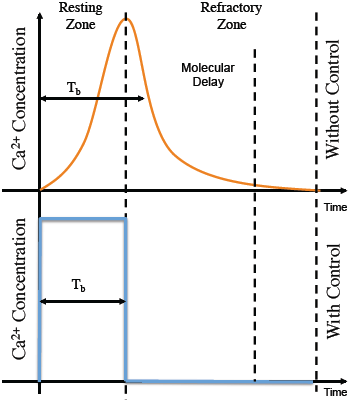
Elimination of the refractory period of a transmitted signal using the proposed control function.

To show the benefits of the proposed control method for molecular communication, a data rate analysis is presented of a single hop Ca^2+^ molecular communication system using astrocytes. The data rate in such system can be computed using: 1*/*(*Tb*∗*Nb*), where *Nb* is the number of bits transmitted. Three *Tb* values were used (5*s*, 10*s*, 50*s*) and compared with the performance for the case when the system integrates the control model as well as without, and varying *Nb* in the process. As demonstrated in Fig. 14 the elimination of the refractory time provides substantial benefit for improving the data rate for all the *Tb* values. On average, the refractory time takes up to five seconds to be completed. This time is needed for a full oscillation cycle from the signalling process of the cell. However, such oscillatory process can also be eliminated with the proposed feed-forward and feedback control technique achieving higher data rate values. For example, with the highest values of data rate (*Tb* = 5*s*), without control reaches a limit of 1000bps while with control reaches 2000 bps.

**Fig. 14:**
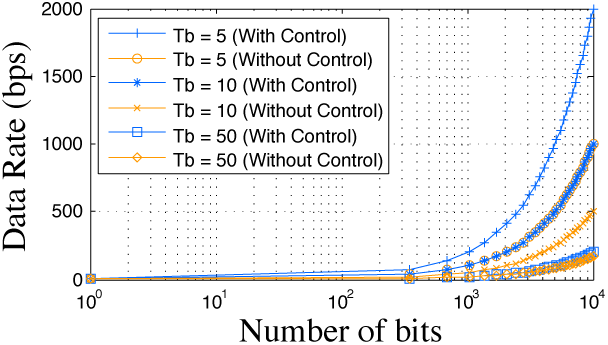
Data rate performance of a Ca^2+^- signalling molecular communication system with *Tb* values (5*s*, 10*s*, 50*s*). Comparison of the performance when the *feedforward and feedback* control technique is applied against the system with no control technique that has to wait for longer refractory periods.

*5) Molecular Gain:* Due to the equal diffusion direction probability (Eq. 27), Ca^2+^ may not reach the *Rx*. Naturally, the amplitude of the received signal is negatively affected by longer distances between the *Tx* and *Rx*. This phenomenon is analysed using gain, which is calculated using the following formula [27]:

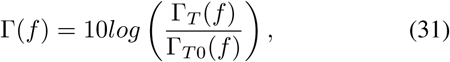

where Γ_*T*_(*f*) is the average peak concentration and Γ_*T*0_(*f*) is the initial peak concentration.

Ideal control of the received Ca^2+^ ions leads to the maintenance of gain performance over distance, Fig. 15. The equaliser provides total recovery in both transmitter and receiver. However, these results might be different for other modulation schemes different than the OOK. On top of that, there must be a distance limit, where signals cannot be reached, and the activation of the control technique cannot be performed.

**Fig. 15:**
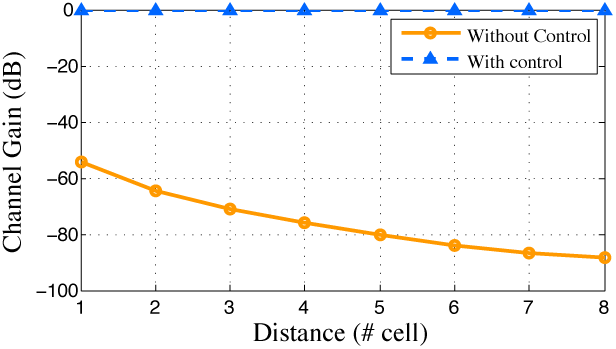
Gain performance of a Ca^2+^- signalling molecular communication system with T*b* values (5*s*, 10*s*, 50*s*). Comparison the performance when the *feedforward and feedback* control technique is applied against the system with no control technique that has to wait for longer refractory periods.

*6) Molecular Delay:* To fairly measure the amount of molecules that are transmitted as well as the time it travels over a certain distance of cellular tissue, the molecular delay proposed in [27] is used to capture this phenomenon. In this way, delay is appropriately measured for each bit, since the information is encoded into a molecular concentration of Ca^2+^ signals. The proposed method is modelled by the following formula:

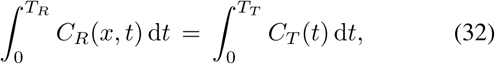

in which, *C*_*R*_ is a function that returns the Ca^2+^ concentration of a *Rx* at time *t*, *C*_*T*_ returns the *Tx* Ca^2+^ concentration at time *t*, *T*_*T*_ is the time slot length of the *Tx* and *T*_*R*_ is the time slot length of the *Rx*. Here, the end-to-end delay will be *T*_*R*_.

Delay is considerably reduced due to the contribution of the regenerated molecules in the receiver by the feed-forward feedback control technique in the form of the equaliser, Fig. 16. The elimination of the refractory periods is another contributor to the delay performance since the remaining interbit interference is also minimised. The difference of delay compared to gain performance is that delay performance should be similar even with different modulation schemes that keep the signal energy levels the same. This result is significant for the minimization of distance effects over the delay, decreasing around an average of 60% delay values per unit distance (number of cells away).

**Fig. 16:**
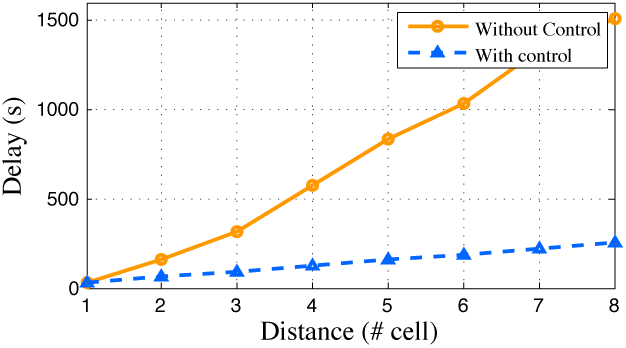
Delay performance of a Ca^2+^- signalling molecular communication system with T*b* values (5*s*, 10*s*, 50*s*). Comparison of the performance when the *feedforward and feedback* control technique is applied against the system with no control technique that has to wait for longer refractory periods.

## VIII. Discussion

In this section, two main applications are explored for the proposed Ca^2+^ control method, including prevention of neurodegenerative diseases and speeding up molecular communication. A significant impact on these topics will result from the utilisation of the proposed method.

### A. Prevention of Neurodegenerative Diseases

Approximately 24 million people worldwide have dementia or neurodegenerative diseases with an annual cost of about $226 billion in the U.S. alone [36], [37]. Causes of such remain unknown, and only conventional symptomatic treatments are available for improving the patients’ health. This is achieved with drugs that target the symptoms alone, neither treating the underlying disease or at best delaying its progression. However, as they progress, brain cells die, and connections among cells are lost, causing the disease symptoms to worsen [38].

A major issue with drug delivery techniques to the brain is the numerous protective barriers that encapsulate the central nervous system, and one of this example is the bloodbrain barrier [38]. Overcoming the blood-brain barrier can be achieved through biotechnology, synthetic biology, as well as nanotechnology, and this can lead to efficient and directed therapeutic tools. The control model proposed in this paper can be developed from a combination of nanoparticles and gene therapy that are used to control the Ca^2+^ signalling, as well as synthetic biology, towards future *in-vivo* settings [17]. Novel self-assembly nanoparticles can bypass the blood-brain barrier and stimulate astrocytes based on existing Ca^2+^ and IP_3_ values. In the case of synthetic biology, programming of cellular signalling pathways can be achieved that can also lead to stable levels of Ca^2+^ ions directly, preventing fluctuations that may result in diseases. This is a more complex solution compared to the one proposed in this paper. Some works have investigated mechanisms to use synthetic biology to engineer neurons to achieve sensitivity to light at a particular wavelength, providing a new alternative for externally controlled neural stimulation [39].

### B. Molecular Communication

Low data rates are a natural characteristic of molecular communication systems due to many factors including stochastic propagation delay as well as excessive noise in the environments [40]. These are due to the natural biological processes that result in poor communication performance. The contribution provided in this paper is far from solving the problem of integrating control theory and synthetic biology with molecular communication scenarios due to the plurality of biological communication channels that are very specific. Therefore, more integration work of engineered processes will be required to counter these natural processes that can affect the performance. This engineering process will usually come through integration of nanotechnology through components and materials to control the biological process or through manipulation of cell using techniques from synthetic biology.

In Section VII-4, the proposed control technique is beneficial to limit the refractory period to increase the overall performance including data rate, delay and gain improvements. However, the proposed control approach is not just limited to that, but also can be extended to perform other communication functionalities that will improve modulation and noise cancellation. Recent work has proposed a modulation technique for a Ca^2+^-signalling based molecular communication system [41], where digital modulation schemes such as *On-Off Shift-Keying* (*OOK*) were used in conjunction with various error control techniques. In [26], noise in Ca^2+^-signalling molecular communication was studied and quantified showing that its high concentration can negatively affect the communication system performance. The noise will emanate at the transmitter as the Ca^2+^ waves are stimulated, along the path as the waves are propagated, and at the receiver, as they stimulate Ca^2+^ ions to receive digital bits. Therefore, based on this scenario, and also on our technique by which the effect of noise can be limited, the control model can be engineered into the cells of the tissue to cancel noise as they propagate along the channel. This can be achieved by restricting the quantity of IP_3_ for each cell that represents the transmitter, along the propagation path, as well as the receiver.

## IX. Conclusions

In recent years, nanotechnology has brought together some different disciplines including synthetic biology and engineering, where the objective is to develop novel health care solutions to detect, prevent and cure diseases. This includes the field of molecular communication, where its aim is to model and construct biological communication systems for inter and intracellular cellular signalling. This new area of research seeks to develop new approaches for detecting and preventing diseases that can emerge from impairments in the communication process, as well as create artificial communication processes that connect a network of nanomachines. This paper investigated one particular type of molecular communication that utilises Ca^2+^ signalling between astrocyte cells and pre and post-synaptic neurons. This three-way communication process is known as the tripartite synapse. In particular, the focus is on the application of a feed-forward and feedback control technique to maintain the stability of Ca^2+^ levels as intercellular signalling is conducted between the cells. The application of the control model achieved firstly, a tight regulation of Ca^2+^ concentration, demonstrably maintaining a stable level in order to minimise any fluctuations, and secondly, an improvement of the overall performance in molecular communication using Ca^2+^-signalling in astrocyte cells. Previous studies have shown that Ca^2+^-signalling in cellular tissue can lead to a significant quantity of noise within the environment, impairing the overall system performance. However, applying the control model has resulted in the reduction in the refractory period of the Ca^2+^-signalling leading to smaller time-slots for bit transmissions, and higher data rates. The control model proposed in this paper can pave the way for novel techniques for disease prevention, as well as mechanisms to improve the performance of molecular communication systems. Furthermore, future work be carried out to investigate improvements of neural activity by enhancing glutamate propagation through control of astrocyte Ca^2+^ signalling.

---

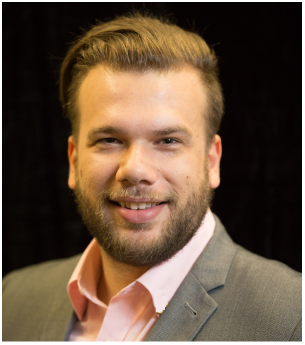

**Michael Taynnan Barros** was born in Campina Grande, Brazil, 1990. He is currently an Irish Research Council Government of Ireland Postdoctoral research fellow associated with the TSSG, WIT. Michael received his PhD in Telecommunication Software at the Waterford Institute of Technology in 2016, M.Sc. degree in Computer Science at the Federal University of Campina Grande in 2012 and B.Tech. degree in Telematics at the Federal Institute of Education, Science and Technology of Paraiba in 2011. He has published over 40 research papers in diverse journals such as IEEE Transactions on Communications, IEEE Transactions on Nanotechnology, and conferences in the area of wireless communications, optical communications, ad-hoc networks, as well as molecular and nanoscale communications. He is also a reviewer for many journals and participated as technical program committee and reviewer for various international conferences. In 2017, he served as the be the Technical Program Co-chair for the 3rd International Workshop on Nanoscale Computing and Communications (NsCC) held in conjunction with NEW2AN conference, the chair of the 5GPPP Network Management, QoS and Security Working Group and the Chair of the 2nd Network Management, QoS and Security for 5G Networks held in conjunction with the EuCNC. Interests in Molecular Communications, Nanonetworks and 5G Technology for Connected Health.

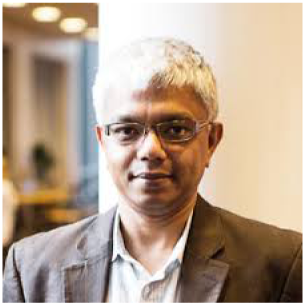

**Subhrakanti Dey** (M’96 - SM’06) was born in India in 1968. He received the B.Tech. and M.Tech. degrees from the Department of Electronics and Electrical Communication Engineering, Indian Institute of Technology, Kharagpur, in 1991 and 1993, respectively, and the Ph.D. degree from the Department of Systems Engineering, Research School of Information Sciences and Engineering, Australian National University, Canberra, in 1996. e is currently a Professor with the Institute of Telecommunications Research (ITR), University of South Australia, Adelaide. Prior to this, he was the Professor of Wireless Sensor Networks with Dept of Engineering Sciences in Uppsala University, Sweden during 2013-2017, and a Professor with the Department of Electrical and Electronic Engineering, University of Melbourne, Parkville, Australia, from 2000 until early 2013. From September 1995 to September 1997, and September 1998 to February 2000, he was a Postdoctoral Research Fellow with the Department of Systems Engineering, Australian National University. From September 1997 to September 1998, he was a Postdoctoral Research Associate with the Institute for Systems Research, University of Maryland, College Park. His current research interests include networked control systems, wireless communications and networks, signal processing for sensor networks, and stochastic and adaptive estimation and control. Professor Dey currently serves on the Editorial Board of the IEEE TRANSACTIONS ON SIGNAL PROCESSING and Elsevier SYSTEMS AND CONTROL LETTERS. He was also an Associate Editor for the IEEE TRANSACTIONS ON SIGNAL PROCESSING during 2007-2010 and the IEEE TRANSACIONS ON AUTOMATIC CONTROL during 2004-2007

## Footnotes

1 One type of gliotransmitter, which activates certain signalling processes in the cells.

## References

[1] B. Atakan, S. Balasubramaniam, and O. B. Akan, “Body area nanonetworks with molecular communications in nanomedicine,” IEEE Communications Magazine, vol. 50, pp. 28–34, 2012.

[2] L. Felicetti, M. Femminella, G. Reali, and P. Liò, “Applications of molecular communications to medicine: A survey,” Nano Communication Networks, vol. 7, pp. 27–45, 2016.

[3] I. Akyildiz, M. Pierobon, S. Balasubramaniam, and Y. Koucheryavy, “The internet of bio-nano things,” IEEE Communications Magazine, vol. 53, no. 3, pp. 32–40, March 2015.

[4] S. A. Wirdatmadja, M. T. Barros, Y. Koucheryavy, J. M. Jornet, and S. Balasubramaniam, “Wireless optogenetic nanonetworks: Device model and charging protocols,” arXiv preprint arXiv:1706.06495, 2017.

[5] I. Akyildiz, F. Fekri, R. S. C. R. Forest, and B. K. Hammer, “Monaco: fundamentals of molecular nano-communication networks,” IEEE Wireless Communications, vol. 19, no. 5, pp. 12–18, 2012.

[6] M. Pierobon and I. F. Akyildiz, “Capacity of a diffusion-based molecular communication system with channel memory and molecular noise,” IEEE Transactions on Information Theory, vol. 59, no. 2, pp. 942–954, 2013.

[7] F. Mesiti, M. Veletc, P. A. Floor, and I. Balasingham, “Astrocyte-neuron communication as cascate of equivalent circuits,” Nano Communication Networks, 2015.

[8] E. A. Newman, “New roles for astrocytes: Regulation of synaptic transmission,” Trends in Neuronscience, vol. 36, no. 10, pp. 536–542, 2003.

[9] S. Nadkarni and P. Jung, “Dressed neurons: modeling neural-glial interactions,” Physical Biology, vol. 1, no. 1, p. 35, 2004.

[10] Q.-S. Liu, Q. Xu, J. Kang, and M. Nedergaard, “Astrocyte activation of presynaptic metabotropic glutamate receptors modulates hippocampal inhibitory synaptic transmission,” Neuron Glia Biology, vol. 1, pp. 307–316, 11 2004.

[11] C. Luscher and R. C. Malenka, “Nmda receptor-dependent long-term potentiation and long-term depression (ltp/ltd),” Cold Spring Harbor Perspectives in Biology, vol. 4, no. 6, p. a005710, 2012.

[12] M. M. Halassa, T. Fellin, and P. G. Haydon, “The tripartite synapse: roles for gliotransmission in health and disease,” Trends in Molecular Medicine, vol. 13, no. 2, pp. 54–63, 2007.

[13] G. Perea, M. Navarrete, and A. Araque, “Tripartite synapses: astrocytes process and control synaptic information,” Trends in Neurosciences, vol. 32, no. 8, pp. 421–431, 2009.

[14] P. Raul, S. Manyam, P. Pagilla, and S. Darbha, “Output regulation of nonlinear systems with application to roll-to-roll manufacturing systems,” IEEE/ASME Transactions on Mechatronics, vol. 20, no. 3, June 2015.

[15] K. Le Meur, J. Mendizabal-Zubiaga, P. Grandes, and E. Audinat, “Gaba release by hippocampal astrocytes,” Frontiers in computational neuroscience, vol. 6, 2012.

[16] M. Pitta, M. Goldberg, V. Volman, H. Berry, and E. Ben-Jacob, “Glutamate regulation of calcium and ip3 oscillating and pulsating dynamics in astrocytes,” Journal of Biological Physics, vol. 35, pp. 383–411, 2009.

[17] M. T. Barros and S. Dey, “Set point regulation of astrocyte intracellular ca2+ signalling,” in The 17th IEEE International Conference on Nanotechnology (IEEE NANO 2017), 2017, pp. 315–320.

[18] J. S. Park, B. Rhau, A. Hermann, K. A. McNally, C. Zhou, D. Gong, O. D. Weiner, B. R. Conklin, J. Onuffer, and W. A. Lim, “Synthetic control of mammalian-cell motility by engineering chemotaxis to an orthogonal bioinert chemical signal,” Proceedings of the National Academy of Science, vol. 111, pp. 5896–5901, 2014.

[19] J. E. Toettcher, O. D. Weiner, and W. A. Lim, “Using optogenetics to interrogate the dynamic control of signal transmission by the ras/erk module,” Cell, vol. 155, no. 6, pp. 1422–1434, 2003.

[20] K. C. Heyde and W. C. Ruder, “A model of a synthetic biological communication interface between mammalian cells and mechatronic systems,” IEEE Transactions on NanoBioscience, vol. 15, no. 8, pp. 864–870, 2016.

[21] S. L. Zhang, A. V. Yeromin, X. H. F. Zhang, Y. Yu, O. Safrina, A. Penna, J. Roos, K. A. Stauderman, and M. D. Cahalan, “Genome-wide rnai screen of ca2+ infiux identifies genes that regulate ca2+ release-activated ca2+ channel activity,” Proceedings of the National Academy of Science, vol. 103, pp. 9357–9362, 2006.

[22] H. Ye, M. D. E. Baba, R. W. Peng, and M. Fussenegger, “A synthetic optogenetic transcription device enhances blood-glucose homeostasis in mice,” Science, vol. 332, pp. 1565–1568, 2011.

[23] T. S. Gardner, C. R. Cantor, and J. J. Collins, “Construction of a genetic toggle switch in escherichia coli,” Nature, vol. 403, no. 6767, pp. 339–342, 2000.

[24] A. Gerbino, W. C. Ruder, S. Curci, T. Pozzan, M. Zaccolo, and A. M. Hofer, “Termination of camp signals by ca2+ and gai via extracellular ca2+ sensors,” Journal of Cell Biology, vol. 171, pp. 303–312, 2005.

[25] B. W. Lau, M. Colella, W. C. Ruder, M. Ranieri, S. Curci, and A. M. Hofer, “Deoxycholic acid activates protein kinase c and phospholipase c via increased ca2+ entry at plasma membrane,” Gastroenterology, vol. 128, no. 3, pp. 695–707, 2005.

[26] M. T. Barros, S. Balasubramaniam, B. Jennings, and Y. Koucheryavy, “Transmission protocols for calcium signaling based molecular communications in deformable cellular tissues,” IEEE Transactions on Nanotechnology, vol. 13, no. 4, pp. 779–788, 2014.

[27] M. T. Barros, S. Balasubramaniam, and B. Jennings, “Comparative endto-end analysis of ca2+ signaling-based molecular communication in biological tissues,” IEEE Transactions on Communications, vol. 63, no. 12, pp. 5128–5142, December 2015.

[28] M. T. Barros, S. Balasubramaniam, and B. Jennings “Using information metrics and molecular communication to detect cellular tissue deformation,” IEEE Transactions on Nanobioscience, vol. 13, no. 3, pp. 278–288, 2014.

[29] J. Lallouette, M. D. Pitta, E. Ben-Jacob, and H. Berry, “Sparse short-distance connection enhance calcium wave propagation in a 3D model of astrocytes networks,” Frontiers in Computation Neuroscience, vol. 8, no. 45, pp. 1–18, 2014.

[30] D. T. Gillespie, “Exact stochastic simulation of coupled chemical reactions,” Journal of Physical Chemistry, vol. 81, no. 25, pp. 2340–2361, 1977.

[31] T. Nakano and J.-Q. Liu, “Design and analysis of molecular relay channels: An information theoretic approach,” IEEE Transactions on NanoBioscience, vol. 9, pp. 213–221, 2010.

[32] R. Janicek, M. Hotka, J. A. Zahradnikova, A. Zahradnikova, and I. Zahradnak, “Quantitative analysis of calcium spikes in noisy fiuorescent background,” PLoS ONE, vol. 8, no. 5, p. e64394, 05 2013.

[33] A. Gaur, A. Midha, and A. L. Mbatia, “Applications of nanotechnology in medical sciences,” Asian Journal of Pharmaceutical Sciences, vol. 2, pp. 80–85, 2008.

[34] S. Orrenius and P. Nicotera, “The calcium ion and cell death,” Journal of Neural Transmission. Supplementa, vol. 43, pp. 1–11, 1994.

[35] F. M. LaFerla, “Calcium dyshomeostasis and intracellular signalling in alzheimer's disease,” Nature Reviews Neuroscience, vol. 3, pp. 862–872, 2002.

[36] A. Association, “Alzheimer's disease facts and figures.” [Online]. Available: https://www.alz.org/facts/downloads/facts_figures_2015.pdf

[37] C. P. Ferri, M. Prince, C. Brayne, and et al., “Global prevalence of dementia: A delphi consensus study,” Lancet, vol. 366, pp. 2112–2117, 2005.

[38] G. Modi, V. Pillay, and Y. E. Choonara, “Advances in the treatment of neurodegenerative disorders employing nanotechnology,” Annals of the New York Academy of Sciences, vol. 1184, pp. 154–172, 2010.

[39] S. A. Wirdatmadja, S. Balasubramaniam, Y. Koucheryavy, and J. M. Jornet, “Wireless optogenetic neural dust for deep brain stimulation,” in 2016 IEEE 18th International Conference on e-Health Networking, Applications and Services (Healthcom), Sept 2016, pp. 1–6.

[40] M. T. Barros, “Ca2+-signaling-based molecular communication systems: Design and future research directions,” Nano Communication Networks, vol. 11, no. 1, pp. 103–113, 2017.

[41] M. T. Barros, S. Balasubramaniam, and B. Jennings, “Error control for calcium signaling based molecular communications,” in 47th Annual Asilomar Conference on Signals, Systems, and Computers, 2013.

